# A Comparison of Mechanisms Driving Lesion Outcomes during Lung Tumor and Tuberculosis Granuloma Formation

**DOI:** 10.64898/2026.02.25.708029

**Authors:** Christian T. Michael, Maral Budak, Denise Kirschner

## Abstract

Small cell lung cancer (SCLC) and tuberculosis (TB) are both deadly diseases that present with spatially complex lung lesions. These lesions share many similarities, including several key spatial interactions between T cells and macrophages. Both SCLC and TB present with significant heterogeneity, both in terms of progression of disease and responses to treatment; current experimental methods have few tools to investigate the spatiotemporal evolution of these lesions within human lungs. We have applied our computational agent-based model, *GranSim*, to extensively study heterogeneity of TB granuloma scale formation, infection outcome and treatment in detail. We introduce *TumorSim*, an analogous agent-based model designed to understand the heterogeneity of SCLC lung tumors. *TumorSim* mechanistically and spatio-temporally captures immune-tumor interactions, many of which are well-studied in isolation, including cytokine-based recruitment of adaptive cells and PD1/PDL1-based inhibition of cytotoxic T-cell activity. Drawing from known lung immunology as well as literature on lung tumor responses, we define and explore a wide set of parameters to characterize *TumorSim* behavior using global sensitivity analysis. We compare factors that drive dynamics of both SCLC tumors and TB granulomas. As model validation, sensitivity analysis captures several well-known correlates of improved SCLC outcomes including macrophage-mediated cytotoxic T-cell recruitment. Surprisingly, both models predict a two-phase formation process occurring with an abrupt change in tumor/granuloma dynamics upon arrival of adaptive immune cells into the lung from lung-draining lymph nodes. Simulations suggest that while CCL5 is associated with improved tumor control later during tumor growth, CCL5 plays a pro-tumor role early during tumor growth by recruiting regulatory T cells. We also find that, similar to virtual TB granulomas, *TumorSim* tumors are increased in volume when immunosuppressive mechanisms outweigh pro-inflammatory responses. This novel tumor model can serve as a basis for future studies on lung tumor-immune dynamics to study both immunotherapeutics and anti-cancer drugs.

## 1. Introduction

Small cell lung cancer (SCLC) and pulmonary Tuberculosis (TB), two leading causes of death worldwide, both manifest in lungs with lesions that share anatomic and functional similarities that directly impact disease progression and responses to therapies. SCLC, an aggressive form of lung cancer with a <15% two-year survival, accounts for over 10% of lung cancer cases. TB is a bacterial disease caused by infection with *Mycobacterium tuberculosis* (Mtb), and is the world’s leading cause of death by single infectious pathogen and has been for decades^1^. Discrete lung masses form in and on lung stroma during both SCLC and pulmonary Mtb infection, and in each disease these masses result from recruitment of multiple, overlapping immune cells to both lesions and their ensuing complex interactions. In fact, due to the similarities between TB and SCLC, TB patients are often misdiagnosed with lung cancer based on PET-CT scans in regions with high TB incidence^2,3^. Moreover, many studies have provided evidence that patients with TB have an increased risk of lung cancer^4-7^. Treatment of both SCLC and TB presents formidable challenges necessitating multi-drug regimens designed to overcome heterogeneous spatial drug delivery and pre-existing and/or acquired drug resistance. While effective (though intensive and expensive) treatment for TB exists^8^, current therapies for SCLC fail for many patients^9,10^ emphasizing the need for us to better understand basic immunological mechanisms and drivers of tumor heterogeneity. A mechanistic computational model can help identify these factors.

Lung cancer, especially SCLC, has many features in common with TB. TB granulomas and lung cancer lesions pose remarkably similar challenges to disease elimination via immune response or treatment (Figure 1). In both, an effective immune response is impaired in the lung microenvironment due to increased immunosuppressive cells (myeloid-derived suppressor cells or regulatory T cells) or T-cell exhaustion^11-13^. Spatially-heterogeneous architectures of granulomas and tumors, including poorly developed vasculature and distorted extracellular matrix, impair drug delivery and T-cell infiltration into both types of lesions^14-16^. The resulting heterogeneous distribution of drugs throughout lesions contributes towards the commonly-observed outcomes of drug resistance and subsequent treatment failure, especially in combination with pre-existing or acquired mutations in granulomas/tumors due to nutrient-poor lesion microenvironments^17,18^. These challenges necessitate extended courses of treatments for both diseases with multiple drugs^19,20^.

**Figure 1:**
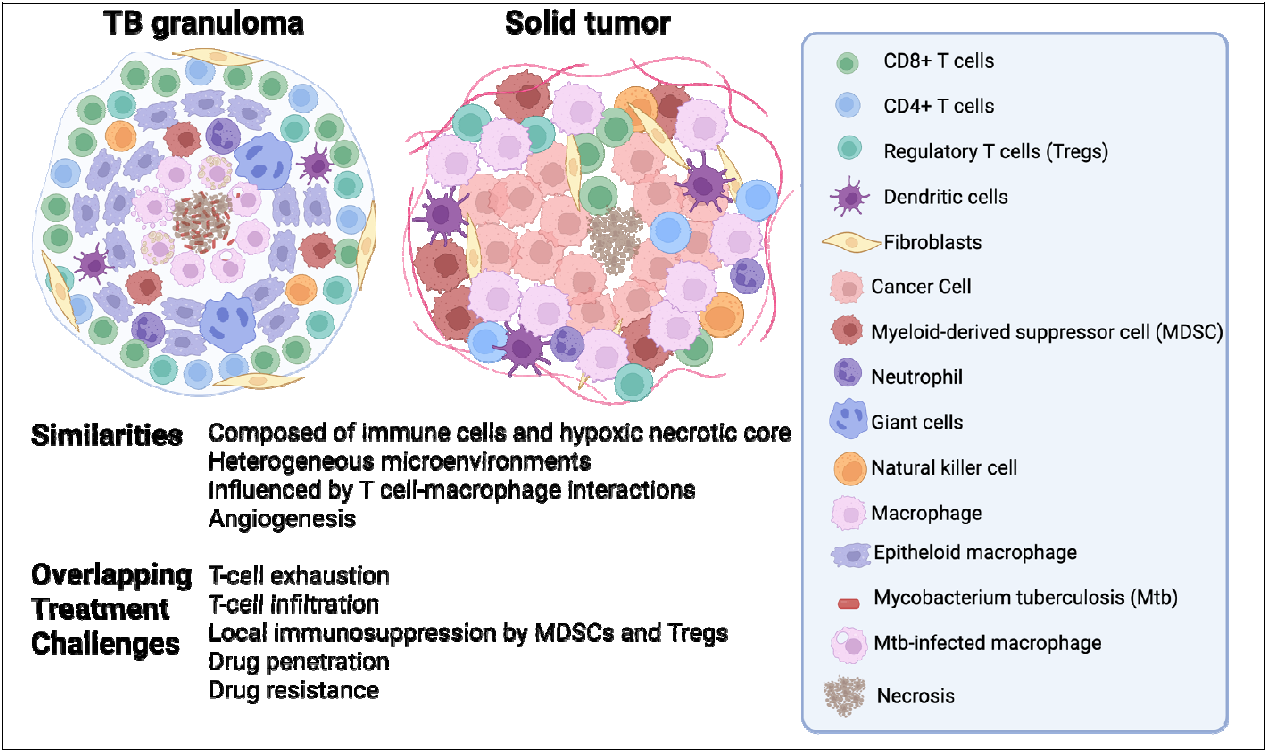
Summary comparison of tuberculosis (TB) granuloma and solid tumor lesion microenvironments. Both lesions comprise multiple types of immune cells clustering around Mtb infection and neoplasm, respectively. Initial interaction between macrophages and dendritic cells result in the recruitment of an adaptive immune response. Activity of apoptosis-inducing leukocytes (cytotoxic T cells, natural killer cells, pro-inflammatory macrophages) is critical for effective resolution and is hampered by anti-inflammatory leukocytes such as regulatory T cells, anti-inflammatory macrophages, and myeloid-derived suppressor cells^4,9^. The core of each lesion is often hypoxic and necrotic^9,26^. Created in BioRender. Michael, C. (2026) https://BioRender.com/1a1nvb3

TB lung granulomas and lung tumors share anatomic and functional similarities that directly impact disease progression and response to therapy^4^. In response to infection with Mtb or formation of cancer cells, the host recruits a wide variety of immune cells, including effector and immunosuppressive myeloid lineage cells and T lymphocytes, and other stromal cell types such as activated fibroblasts. This results in the emergence of complex, and spatially heterogeneous structures called granulomas (TB) or solid tumors (cancer) (Figure 1)^11,21-23^. Host immune responses commonly do not eliminate Mtb or cancer cells, producing two outcomes: 1) progressive disease in either case; or 2) clinical latency (TB) or dormancy (cancer), in which a small number of bacteria or cancer cells remain in a viable, growth-arrested or slowly-dividing state for extended periods of time^24,25^. The course of TB and lung cancer is heavily dependent on the interactions between T lymphocytes and macrophages in lesion microenvironments^11,21^.

Emergent lung lesions such as tumors and granulomas are challenging to study, as gross predictive power can be elusive despite knowing the myriad of constituent mechanisms and key cell types present. An effective method for identifying principle drivers of clinical outcomes are *in silico*, mechanistic, multivariate, mathematical models. When calibrated and validated to experimental and clinical datasets, multi-scale models are useful and cost-efficient at capturing complexities of a biological system that may be challenging to probe by experimental methods. We have previously developed a multi-scale mechanistic hybrid agent-based model, *GranSim*, that recapitulates heterogeneous outcomes of lung granuloma progressions, entirely emergent from cell-scale immune responses to pulmonary Mtb infection^27-30^. In this work, we create a novel, tandem hybrid agent-based model to understand factors that contribute to lung solid tumor formation and heterogeneity. We introduce a new multi-scale model, *TumorSim*, that captures mechanisms driving SCLC lung tumor formation dynamics. We compare and contrast *TumorSim* to the dynamics of our well-characterized *GranSim* model behavior to investigate similarities and differences between mechanisms of TB and SCLC. Our goal is to harness cross-talk between these two disparate disciplines (infection disease and cancer) that will help foster new ideas and research advancement for both.

## 2. Methods

To study emergent lesion structures that arise such as SCLC tumors or TB granulomas, we need to explicitly represent cell-and molecular scale details. Several simplified models of tumors exist, though these models focus on large-scale measurements and embed mechanistic hypotheses within the structure their governing equations^31^. We choose to model both the temporal and spatial aspects of SCLC lung tumors as that have arisen as a key feature in treatment of lung diseases^9,32-35^. Agent-based models (ABMs) represent systems biology systems as collections of interacting *agents* (e.g., cells) that interact to form emergent structures; direct representation allows for mechanistic probing of how smaller-scale agents affect larger-scale outcomes (e.g., tumors and granulomas). ABMs allow for explicit representation of cell-scale mechanism and modular updates as new information become available and have been used for a variety of applications^36-38^. Published cancer models have been formulated as ABMs to address important questions around immune dynamics^39-46^, growth of metastases^47-51^, drug resistance^52-54^, tumor immunotherapy, tumor-enhancing immune/stromal cells, and tumor heterogeneity^55^. Gong et al.^56^ developed a multiscale, three-dimensional (3D) ABM of interactions between immune and cancer cells, which revealed emergent global behaviors during tumor development and therapy. Menezes et al. and Calopiz et al. created ABMs detailing the pharmacokinetics of antibody drug conjugate treatment of solid tumors, relating drug penetration to overall impact on treatment efficacy for established tumors^57,58^.

Basing the framework we have created and curated for over 20 years for an ABM capturing TB granulomas, *GranSim, we* formulate *TumorSim* as an ABM representing SCLC tumors. We have previously used *GranSim* to study immune and bacterial dynamics during infection^27,30,59-62^, antibiotic treatment^63-69^, perform virtual clinical trials to predict optimal treatment strategies^63-67,69-72^, also incorporating drug resistance^73^. *GranSim* reproduced temporal dynamics of cytotoxic T cells and cancer cells during tumor progression, as well as spatial distributions of these cells. As SCLC tumors share many fundamental components, we create *TumorSim* on our foundational *GranSim* computational architecture platform. Here, we create novel agents, rules and features (Hypoxia inducible factor 1-α (HIF1α), oxygen, interferon gamma (IFN□), cancer cells etc.) and use wide ranges of parameters to explore range of tumor model outputs. A comparison between mechanisms included in *GranSim* and *TumorSim* is shown in Table 1 showing both overlap and distinction between features.

**Table 1:** Comparison between *GranSim* and *TumorSim* mechanisms.

| Table 1: Comparison between *GranSim* and *TumorSim* mechanisms |  |  |
| --- | --- | --- |
|  | *GranSim* | *TumorSim* |
| Initialization | - Resting macrophages and vascular sources randomly distributed on simulation grid<br>- One macrophage infected with one mycobacterium at the center of the simulation grid | - Vascular sources randomly distributed on simulation grid<br>- A circular neoplasm of ten cancer cell diameter at the center of the simulation grid<br>- Few, and M1 macrophages at the center of the simulation grid |
| Macrophages | - Present as resting (Mr), activated (Ma), infected (Mi), chronically infected (Mci)<br>- Ma kills Mtb effectively<br>- Mi provides a permissive niche for Mtb to grow<br>- Mci cannot kill Mtb and bursts. | - Present as M1 and M2<br>- M1 kills cancer cells<br>- M2 deactivates Tcyt<br>- IFNγ polarizes M2 toward M1 phenotype<br>- HIF1α polarizes M1 toward M2 phenotype |
| T cells | - Present as cytotoxic (Tcyt), IFNγ secreting (Tgam) and regulatory (Treg)<br>- Tcyt and Tgam are proinflammatory and induce apoptosis in Mi and Mci<br>- Treg is anti-inflammatory and downregulate Tcyt and Tgam | - Present as Treg and Tcyt<br>- Tcyt is proinflammatory and kills cancer cells, can be PDL1+ or PDL1-<br>- Treg is anti-inflammatory and deactivates Tcyts |
| Vascular blockage | - Vascular sources blocked depending on the amount of necrotization and cell density | - Dynamic vascular permeability based on the level of hypoxia |
| Chemokine-cytokine secretion | - Chemokines: CCL2, CCL5 and CXCL9 secreted by Mi, Ma and Mci<br>- Cytokines: TNFα secreted by Tgam and Tcyt; IFNγ secreted by Tgam; IL-10 secreted by macrophages, Tregs, and nominally by Tcyts | - Chemokines: CCL2, CCL5 secreted by cancer cells, and CXCL9 M1 macrophages<br>- Cytokines: IFNγ secreted by M1 and Tcyt;<br>- HIF1α implicitly secreted by all cells in low-oxygen environments |
| Recruitment | - Macrophages and T cells recruited from vascular sources based on the levels of CCL2, CCL5, CXCL9, and TNF-α | - Macrophages and T cells recruited from vascular sources based on the levels of CCL2, CCL5, CXCL9 |
| Proliferation | - Bacteria proliferate with different rates based on the bacterial subpopulation.<br>- Immune cells do not proliferate. | - Cancer cells proliferate<br>- Immune cells may proliferate |
| Age-related cell death | Immune cells only | No age-related cell death |

### 2.1. TumorSim

To simulate emergence of a SCLC tumor, we create *TumorSim*, which captures key cells known to drive basic SCLC immunobiology. We build *TumorSim* as a multiscale, agent-based model that simulates interactions between immune cells, relevant molecules and cancer leading to formation of SCLC tumors on a 2-D simulation grid representing 6mm x 6mm of lung tissue. *TumorSim* contains two mathematically distinct types of agents.

First are discrete agents that move or act based on rules (e.g., cell-cell interactions) at a slow timescale (*ΔT* = 10 minutes). Discrete agents include vascular sources that serve as a source of cellular recruitment and O_2_ levels. All other discrete agents are cellular and include cancer cells that may or may not express the PDL1 “don’t-kill-me” ligand, cytotoxic CD8+ T cells (Tcyts) that may or may not express PD1, regulatory T cells (Treg), and pro- and anti-inflammatory (M1 and M2) macrophages. These are the minimal set of immune cells known to be heavily involved in SCLC lung lesions^9,35^.

Second are continuous agents that are secreted, consumed, or diffuse at a fast timescale (*ΔT* = 6 seconds). Rules governing continuous agents are discretized partial differential equations (PDEs), allowing a slower timescale. Cells secrete or consume molecules, and rule-based cell-cell interactions occur with probabilities described in the following sections. The continuous agents in *TumorSim* include IFN□ representing a pro-inflammatory factor^34,74^; and CCL2, CCL5, and CXCL9 as chemokines that direct cell migration^33-35^. Further, we uniquely track O_2,_ which diffuses through the microenvrionment and is consumed by cellular agents. We use O_2_ to infer a hypoxia marker, HIF1α that directs cells towards immunosuppressive phenotypes^75,76^. We describe rules and equations in detail in the following subsections.

#### 2.1.1. Initialization

Shortly after the initial malignant transformation, nascent tumor cells may be eliminated by the immune system during an *elimination phase*^*77,78*^. The elimination phase is likely short, and known to involve many components that are outside of the scope of this study, including CD4+ T cells and γδ T cells. For this reason, we initiate our virtual SCLC tumor several days after tumor transformation. As our initial state, we represent a small cluster of cancer cells in lung tissue after immune signals from the elimination phase have allowed for macrophages, Tregs, and Tcyts to begin to be recruited^78^.

*TumorSim* tumors begin as small circular lesions with a 10-cell diameter placed upon a 6mm x 6mm 2D simulation grid. The grid is divided into 20 μm x 20 μm compartments, this size allows one macrophage, T cell, or cancer cell to occupy a grid compartment as in previous cancer ABMs^57,58^. The entire lung grid section is initially populated by tissue densities of D_Tcyt_ cytotoxic T cells (Tcyts), D_Treg_ regulatory T cells (Tregs), and macrophages (D_M1_ for M1 and D_M2_ for M2). Cancer cells are placed in a grid compartment in a circle of diameter *d*_Tumor_ centered in the middle of the tissue grid. We also randomly distribute vascular sources onto the 2D grid with a density D_VS_. This is the density we vascularize the lung with for *GranSim* simulations, as the lung tissue is the same^64^. Simulations initialize with both Tcyts and cancer cells as PD1- and PDL1-, respectively, as SCLC is characterized by low levels of these markers^79^. Initial conditions are shown in Table 2.

**Table 2:** Initialization parameters.

| Parameter | Value/Range | Description |
| --- | --- | --- |
| $d_{\text{Tumor}}$ | 10 cancer cells (200μm) | Initial tumor diameter |
| N<sub>Tcyt</sub> | 0 | Initial number of Tcyts |
| D<sub>Treg</sub> | [0.0005, 0.0008] | Initial density of Treg |
| D<sub>M1</sub> | [0.00003, 0.00005] | Initial density of M1 macrophages |
| D<sub>M2</sub> | [0.0007, 0.001] | Initial density of M2 macrophages |
| D<sub>VS</sub> | 0.0144 | Initial density of vascular sources |

#### 2.1.2. T cells, cancer cells, and the PD1/PDL1 axis

The levels of cytotoxic and regulatory T cells associated with tumors, as well as their exposure to pro/anti-inflammatory cytokines, are strongly associated with tumor prognosis^34,74,80^. T cells may express the programmed cell death protein 1 (PD1) as a surface protein, and tumors cells may produce its ligand (PDL1). This receptor-ligand pair act as a natural brake to prevent T cells from killing healthy tissue, but PDL1 expression by cancer cells inhibit the ability of T cells to kill cancer cells^32^. HIF1α, produced by cells during hypoxia, induces expression of PD1 and PDL1 in Tcyt^75^ and cancer cells^81^, respectively. Conversely, interleukin-12 signaling induces expression of the transcription factor T-bet, which both directly represses PD1 transcription and promotes IFN□ production^82-86^. Similarly, interleukin-2 indirectly regulates the expression of PDL1 on cancer cells via Tcyts, which has been modeled previously in detail^56^.

In *TumorSim*, we coarse-grain each of the above interactions as events that stochastically occur during each large-scale timestep. We represent Tcyts, Tregs, and cancer cells as discrete agents in our simulated tissue grid. Secretion of the continuous agents IFN□ and HIF1α are described in sections 2.1.4 and 2.1.7, respectively. Tcyts may be in the discrete states (i) activated or inactive, and (ii) PD1+ or PD1-; and cancer cells may be PDL1+ or PDL1-. Inactive Tcyts may become activated with higher IFN□ or when they are adjacent with (i.e. within the Moore neighborhood of) a cancer cell. An activated Tcyt adjacent to a cancer cell may kill that cancer cell with a probability based on whether the Tcyt is PD1± and/or the cancer cell is PDL1±. Tregs may deactivate adjacent Tcyts. Tcyts can induce PDL1 expression in adjacent cancer cells. PD1+ Tcyts can be deactivated by PDL1+ cancer cells. removing PD1 on the surface of Tcyts that increases with high levels of IFN□. Equations 1-4 and Table 3 parameterize these interactions:

**Table 3:** T-cell parameters. Ranges in *italics* are broad-range estimates to explore model capability. *We established this exploratory range by finding the maximum and minimum IFN□ molecule counts per grid compartment in pilot simulations. †We sample this range using a log-uniform distribution.

| Parameter | Value/Range | Description |
| --- | --- | --- |
| P<sub>actPD1-TcytKillingPDL+Tumor</sub> | *[0.01, 0.1]*<sup>†</sup> | Probability of activated PD1- Tcyt killing adjacent PDL1+ cancer cells |
| P<sub>actPD1+TcytKillingPDL+Tumor</sub> | 0 | Probability of activated PD1+ Tcyt killing adjacent PDL1+ cancer cells |
| P<sub>actPD1-TcytKillingPDL-Tumor</sub> | *[0.1, 1]*<sup>†</sup> | Probability of activated PD1- Tcyt killing adjacent PDL1- cancer cells |
| P<sub>actPD1+TcytKillingPDL-Tumor</sub> | *[0.01, 0.1]*<sup>†</sup> | Probability of activated PD1+ Tcyt killing adjacent PDL1- cancer cells |
| P<sub>TregDeactTcyt</sub> | *[0.1, 0.9]* | Probability of Treg deactivating adjacent Tcyt |
| P<sub>Tcyt_tumorPDL1</sub> | *[0.01, 1]*<sup>†</sup> | Probability of Tcyt inducing PDL1 expression in adjacent cancer cells |
| P<sub>tumorPDL1deactivatingPD1+Tcyt</sub> | *[0.01, 1]*<sup>†</sup> | Probability of PDL1+ cancer cell deactivating adjacent active PD1+ Tcyt |
| P<sub>TcytActivationByTumor</sub> | *[0.01, 1]*<sup>†</sup> | Probability of Tcyt activation adjacent to a cancer cell |
| V<sub>maxTcytActivation</sub> | *[0.1, 0.9]* | Max IFNγ-dependent Tcyt activation probability |
| C50<sub>TcytActivation</sub> | *[0.5, 3]* * | Half-saturation molecule count for IFNγ-dependent Tcyt activation |
| V<sub>maxPD-PD+TcytConversion</sub> | *[0.01, 1]*<sup>†</sup> | Max HIF1α-dependent PD1- → PD1+ Tcyt conversion probability |
| C50<sub>PD-PD+TcytConversion</sub> | *[0.5, 0.9]* | Half-saturation relative concentration for PD1- → PD1+ HIF1α-dependent Tcyt conversion |
| V<sub>maxPD+PD-TcytConversion</sub> | *[0.01, 1]*<sup>†</sup> | Max IFNγ -dependent PD1+ → PD1- Tcyt conversion probability |
| C50<sub>PD+PD-TcytConversion</sub> | *[0.5, 3]* * | Half-saturation relative concentration for PD1+ → PD1- IFNγ-dependent Tcyt conversion |
| V<sub>maxPDL-PDL+TumorConversion</sub> | *[0.01, 1]* | Max HIF1α-dependent PDL1- → PDL1+ cancer cell conversion probability |
| C50<sub>PDL-PDL+TumorConversion</sub> | *[0.5, 0.9]* | Half-saturation relative concentration for PDL1- → PDL1+ HIF1α-dependent cancer cell conversion |

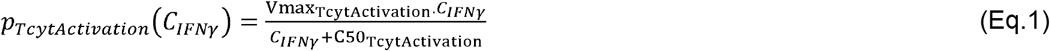

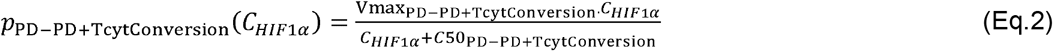

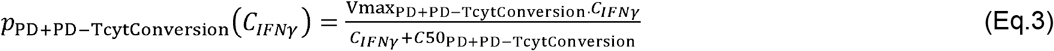

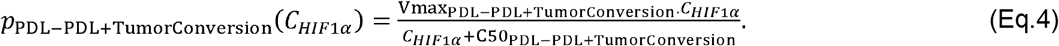

#### 2.1.3. Macrophages

Tumor-associated macrophages are a key component of SCLC and their control^13,33,87^. They can either be pro-inflammatory and help destroy the tumor or be anti-inflammatory and facilitate tumor growth. They also secrete chemokines important to T-cell recruitment (Section 2.1.4). We represent proinflammatory (M1) and anti-inflammatory (M2) macrophages that have been shown to affect tumor progression^88^. M1 macrophages are able to kill cancer cells when adjacent to a cancer cell and M2 macrophages are able to deactivate adjacent Tcyts. M1 can convert to the M2 phenotype in the presence of high HIF1α levels, as products downstream of HIF1α are associated with M2 polarization^89^. M2 can be converted to M1 in the presence of high levels of IFN□ ^90^. Equations 5-6 and Table 4 describe polarization:

**Table 4:** Macrophage parameters. Ranges in *italics* are a broad-range estimate to explore model capability. †We established this exploratory range by finding the maximum and minimum IFN□ molecule counts per grid compartment in pilot simulations.

| Parameter | Value/Range | Description |
| --- | --- | --- |
| $p_{M1killingTumor}$ | [0.1, 0.9] | Probability of M1 killing adjacent cancer cell |
| $p_{M1activatingTcyt}$ | [0.1, 0.9] | Probability of M1 activating adjacent Tcyt |
| $p_{M2deactivatingTcyt}$ | [0.1, 0.9] | Probability of M2 deactivating adjacent Tcyt |
| $V_{max_{M1M2conversion}}$ | [0.1, 0.9] | Max HIF1α-dependent M1→M2 conversion probability |
| $C50_{M1M2conversion}$ | [0.1, 0.9] | Half-saturation parameter for M1→M2 HIF1α-dependent conversion |
| $V_{max_{M2M1conversion}}$ | [0.1, 0.9] | Max IFNγ-dependent M2→M1 conversion probability |
| $C50_{M2M1conversion}$ | [0.5, 3] | Half-saturation molecule count for M2→M1 IFNγ-dependent conversion |

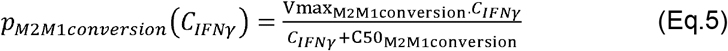

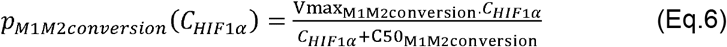

#### 2.1.4. Chemokine-cytokine secretion and chemotaxis

A critical component of immune control of SCLC tumors are cytokines and chemokines that activate immune cells and guide cells toward the developing tumor^35,77^. In *TumorSim*, we represent both IFN□ and several chemokines as continuous variables (units of #molecules/grid compartment) that diffuse through tissue and are eventually cleared through degradation. Proinflammatory cells (M1 macrophages and Tcyts) secrete IFN□ at the fast timestep (6 seconds). IFN□ secretion from Tcyts decreases with higher HIF1α, as downstream products decrease IFN□ secretion from CD8+ T cells (the most common form of Tcyt)^76^. IFN□ secretion by M1 macrophages occurs at a constant rate, though we assume that macrophage production of IFN□ is less than that of Tcyts^74^.

We also assume that cancer cells constitutively produce the chemokines CCL2 and CCL5, known to initially recruit monocytes and T cells, respectively. We also assume that IFN□ can induce M1 macrophages to produce CXCL9, another chemokine associated with Tcyt responses^34,35,91^. Tregs also migrate in response to CXCL9 and CCL5^80,92^. We allow virtual M1 and M2 macrophages to follow CCL2 and CCL5 gradients, though we note this simplification serves to coarse-grain this chemoattraction prior to polarization^93^. There is evidence that SCLC silences CCL2^33^ and CCL5^34^, leading to an affected cytotoxic T-cell response that may undermine PD1/PDL1 blockade treatment, so we vary production rates of CCL2 and CCL5 to study this further. Table 5 and equations 7-8 characterize these interactions:

**Table 5:** IFN□ and chemokine parameters. Ranges in *italics* are broad-range estimates to explore model capability. *Value adapted from *GranSim*.

| Parameter | Value/Range | Description |
| --- | --- | --- |
| $V_{\max_{\text{IFN}\gamma_{\text{Tcyt}}}}$ | *[0.1, 1] (1/s)* | Max IFNγ secretion rate from cytotoxic T cells |
| $C50_{\text{IFN}\gamma_{\text{Tcyt}}}$ | *[0.1, 0.9]* | Half-saturation parameter for HIF1α-dependent IFNγ secretion rate |
| $S_{\text{IFN}\gamma_{\text{M1}}}$ | *[0.01, 0.1] (1/s)* | IFNγ secretion rate from M1 macrophages |
| $D_{\text{IFN}\gamma}$ | 1.4e-8 (cm<sup>2</sup>/s) | Diffusion coefficient of IFNγ |
| $\mu_{\text{IFN}\gamma}$ | 0.00263* (1/s) | Clearance rate of IFNγ |
| $V_{\max_{\text{CXCL9}}}$ | 12* (1/s) | Maximum secretion rate of CXCL9 by M1 macrophages |
| $C50_{\text{CXCL9}}$ | 90 | Half-saturation molecule count for IFNγ-dependent CXCL9 secretion rate by M1 macrophages |
| $S_{\text{CCL2}}$ | *[0, 0.1] (1/s)* | Rate of CCL2 secretion by cancer cells |
| $S_{\text{CCL5}}$ | *[0, 0.1] (1/s)* | Rate of CCL5 secretion by cancer cells |
| $D_{\text{CK}}$ | 5.2e-08 (cm<sup>2</sup>/s) | Diffusion coefficient of CCL2, CCL5, and CXCL9 |
| $\mu_{\text{CK}}$ | 0.000263* (1/s) | Clearance rate of CCL2, CCL5, and CXCL9 |

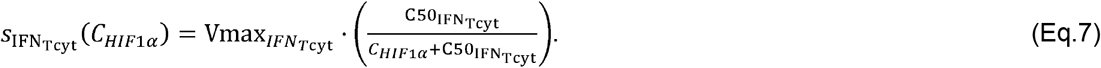

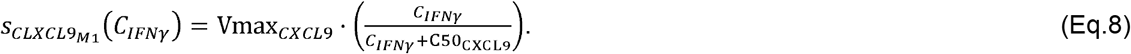

#### 2.1.5. Proliferation

SCLC cancer cells are characterized by rampant proliferation^94^. In *TumorSim*, cancer cells proliferate based on a proliferation time parameter, which determines the time interval needed between two consecutive proliferation events (Table 8). Doubling times for several rapidly-dividing cancer cell types such as glioblastoma and HeLa are between 18-33 hours *in vitro*^95,96^. However, doubling times for SCLC tumors have been measured to be between 25 and 217 days, albeit many such measurements are made by the time the tumors are at the cm-scale^97^. We assume that *in vivo* conditions will prolong doubling time and investigate virtual SCLC cancer cells with doubling times of 36-48 hours. Additionally, some immune cells are known to proliferate in response to cancerous tumor microenvironments^98-100^. We allowed a fraction of immune cells to proliferate based on activation state with cell cycle lengths estimated from literature (as referenced in Table 6). Notably, Tcyts are known to have cell cycle durations as short as 2 hours that slow substantially after several division rounds^100,101^; we choose an 8-hour cell cycle time to account for this eventual slowing. Table 6 characterizes *TumorSim* proliferation.

**Table 6:** Simulated cell cycle durations and frequency of proliferating phenotype by cell type. We estimated cell cycle lengths from the references cited, though true cell cycle durations are dynamically based on many factors that lie beyond the scope of this simulation. Values marked with * are an ansatz, as we could not find specific replication rates in the literature; however, macrophage replication in tumors has been reported^102^.

| Parameter | Value/Range | Description |
| --- | --- | --- |
| $r_{\text{tumor}}$ | [1.5, 2] days | Cancer cell doubling time |
| $p_{\text{tumor}}$ | [0.65, 1] | Probability that cancer cells are proliferating<sup>103</sup> |
| $r_{\text{Treg}}$ | 5 days | Treg<sup>98</sup> proliferation interval |
| $p_{\text{Treg}}$ | 0.01 | Probability that Tregs are proliferating<sup>99</sup> |
| $r_{\text{Tcyt\_rest}}$ | 7.5 days | Inactivated Tcyt<sup>100</sup> proliferation interval |
| $p_{\text{Tcyt\_rest}}$ | 0.03 | Probability that resting Tcyts are proliferating<sup>99</sup> |
| $r_{\text{Tcyt\_act}}$ | 8 hours | Activated Tcyt<sup>100,101</sup> proliferation interval |
| $p_{\text{Tcyt\_act}}$ | 0.05 | Probability that active Tcyts are proliferating<sup>99</sup> |
| $r_{\text{M1}}$ | 1.5 days | M1 macrophage proliferation interval * |
| $r_{\text{M2}}$ | 2 days | M2 macrophage proliferation interval * |
| $p_{\text{M}}$ | [0.05, 0.1] | Probability that M1 or M2 macrophages are proliferating * |

#### 2.1.6. Recruitment

Chemokines associated with inflammation and hypoxia recruit immune cells to the tumor microenvironment^35^. In *TumorSim*, each vascular source has a probability to recruit immune cells based on levels of local chemokines. The chemokines influencing these probabilities are based on identical reasoning to the chemotaxis model^35,93^. We assume that both M1 and M2 macrophages are recruited by CCL2 and CCL5, Tregs are recruited by CCL5, and Tcyts are recruited by both CCL5 and CXCL9. Newly-recruited Tcyts are activated, as we assume activation within lymph nodes occurs prior to their arrival in the tumor microenvironment^104^. Table 7 and equations 8-10 characterize recruitment:

**Table 7:** Recruitment parameters. Ranges are broad-range estimates to explore model capability.

| Parameter | Value/Range | Description |
| --- | --- | --- |
| Rmax<sub>M</sub> | [0.0058, 0.025] | Max recruitment probability of M1 and M2 |
| Rmax<sub>Tcyt</sub> | [0.012, 0.063] | Max recruitment probability of Tcyt |
| Rmax<sub>Treg</sub> | [0.0094, 0.041] | Max recruitment probability of Treg |
| C50<sub>M</sub> | [2.37, 4.62] | Half-saturation molecule count for M1 and M2 chemokine-dependent recruitment |
| C50<sub>Tcyt</sub> | [4.76, 10.27] | Half-saturation molecule count for Tcyt chemokine-dependent recruitment |
| C50<sub>Treg</sub> | [0.94, 2.0] | Half-saturation molecule count for Treg chemokine-dependent recruitment |

**Table 8:**
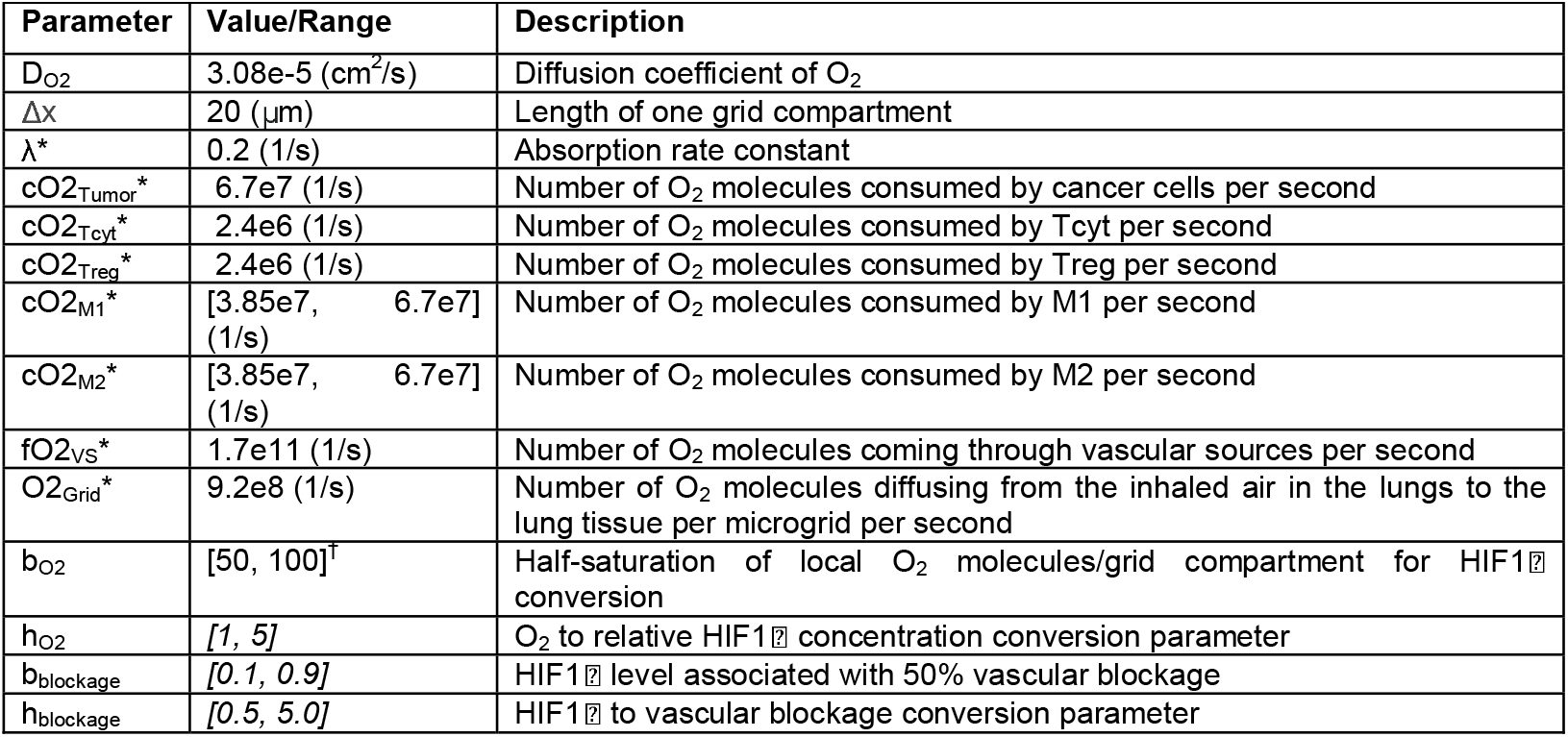
Vascular permeability, O_2_, and HIF1α, and parameters. We present further derivation for parameters marked by * in Supplemental Material 1. †This parameter is based on preliminary simulations where approximately 20mg/L was a common baseline concentration outside of the tumor.

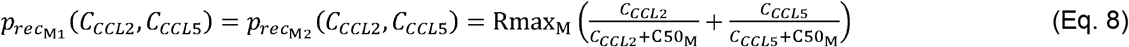

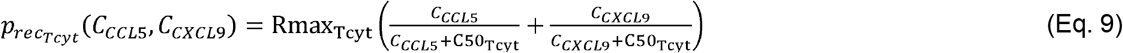

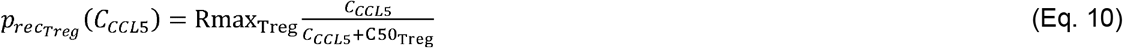

#### 2.1.7 Vascular permeability modeling

Hypoxia, which frequently accompanies solid tumor development, results in vascular blockage that inhibits their ability to supply immune cells to the tumor microenvironment. In *TumorSim*, we model vascular permeability by calculating the blockage for each vascular source, which depends on levels of hypoxia at the source’s location^105^. To assess the level of hypoxia, we calculate O_2_ concentration at steady state within the whole grid using the heat equation and periodic boundary conditions after we initialize an n-by-n grid with the cancer cells, immune cells and vascular sources as

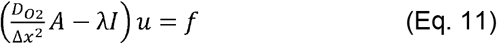

where *D*_*O*2_ is the diffusion coefficient of O2, Δ*x* is the length of one grid compartment, *A* is an n^2^-by-n^2^ Laplacian matrix for periodic boundary conditions, *u* is an n^2^ -by-1 vector of O2 concentrations at the simulation grid and *f* is the n^2^-by-1 / vector for the O2 contribution from sources and sinks (see Table 5 parameter descriptions and values). We define the forcing vector *f* per grid compartment *i* as:

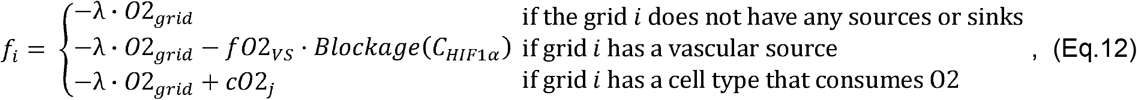

where oxygen consuming cells are Cancer cells, Tcyts, Tregs, M1 macrophages, and M2 macrophages, and *Blockage* (*C*_*HIF*1α_) is defined below (Supplemental Material S1 for additional detail, located at http://malthus.micro.med.umich.edu/TumorSim/).

Hypoxia induces expression of HIF1α^9^. We model relative HIF1α concentration based on steady state O2 concentrations. Relative HIF1α concentration *C*_*HIF*1α_ is a unitless per-grid-compartment value between 0 and 1 that increases with hypoxia (i.e., lower O2 levels), and computed using the following equation (parameterized in Table 8):

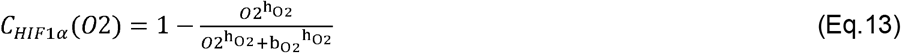

We model vascular blockage based on relative HIF1α (Eq.13, Table 8 for parameterization). We assume vascular sources that are exposed to higher HIF1α are increasingly blocked as higher HIF1α is associated with leaky vascularization via VEGF^106^. The blockage term is between 0 and 1 and is proportional to HIF1α and quantifies the proportion of molecules allowed to pass through each vascular source, where a value of 0 indicates no passage, 0.5 represents 50% passage, etc. (see *f*_i_). Blockage is calculated as:

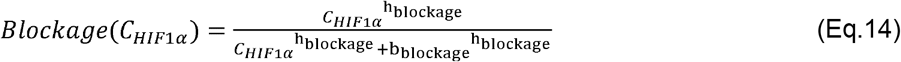

### 2.2. GranSim

The immune cellular composition of lung tumors and granulomas are similar^4^. To compare whether the most influential mechanisms underpinning them are also analogous, we present a brief summary of the components of *GranSim*, which captures key cell types that lead to emergent TB granuloma formation and maintenance. These interactions lead to the emergence of a single lung granuloma that is structurally and functionally representative of non-human primate granulomas, the animal TB model closest to human TB disease and also closely resembles human granulomas^27,30,59-62^.

*GranSim* is a multiscale ABM simulating the interactions between immune cells and bacteria (Mtb) on a 2-D simulation grid representing a 6mm x 6mm lung tissue. Like *TumorSim, GranSim* uses discrete and continuous agents using the same slow and fast timescales (*ΔT* = 10 minutes and *ΔT* = 6 seconds, respectively). Discrete *GranSim* agents include cytotoxic, IFN□ secreting and regulatory T cells; macrophages, Mtb, and vascular sources that serve as loci for cellular recruitment. The grid is divided into 20 μm x 20 μm compartments, we admit a single compartment to contain one macrophage, two T cells, or one macrophage and one T cell. Continuous agents include chemokines (CCL2, CCL5 and CXCL9) and cytokines (TNFα and IFN□, IL-10). Continuous agents are implemented as PDEs, and are updated using an FFT-based PDE solver as in our previous work^107^.

Below is an overview of the major components of *GranSim*. For details and parameter values, see our online documentation at http://malthus.micro.med.umich.edu/GranSim/docs/gransimrules-v2.pdf.

#### 2.2.1. Initialization

TB can be caused by inhaling a single bacterium^108^. As such, we begin with a single macrophage infected with Mtb at the center of the simulation grid. Resting macrophages and vascular sources are randomly distributed onto the 2-D simulation grid, representing otherwise-healthy tissue. We have both slow-timescale (cellular) and fast-timescale (molecular) mechanisms.

#### 2.2.2. Macrophages

Macrophages provide both the main replicative niche for Mtb and once activated, a primary source of bactericidal activity^26^. Macrophages in *GranSim* assume one of four states depending on their activation and infection statuses: resting, activated, infected and chronically infected. Macrophages are in the ‘resting state’ when they are first recruited into the simulation grid. They switch into an ‘activated state’ when they are activated by STAT1 and NFKB-dependent mechanisms and can kill Mtb effectively. They get infected with a defined probability when they are adjacent to Mtb and switch into an ‘infected state’. Infected macrophages provide a permissive niche for Mtb that grow within macrophages. Once the number of Mtb within a macrophage reaches a predefined threshold, then the macrophage switches from the ‘infected state’ into the ‘chronically infected state’, where it cannot clear the Mtb or move due to highly infected status. Once the number of intracellular Mtb reaches the carrying capacity, the chronically infected macrophage bursts and dies.

#### 2.2.3. T cells

Mtb-specific T cells are critical for effective control of Mtb, both through inducing apoptosis of infected macrophages, activation of resting macrophages (allowing them to effectively kill Mtb)^109,110^. Tregs cells are immunosuppressive, and their activity is associated with poor granuloma outcomes^110^. *GranSim* represents three types of T cells: cytotoxic (Tcyt), IFN□-secreting (Tgam), and regulatory (Treg). Cytotoxic T and IFNγ-secreting T cells secrete TNFα, but only IFN□-secreting T cells secrete IFN□. Cytotoxic T cells induce perforin/granulysin-mediated apoptosis, whereas IFN□ -secreting T cells induce Fas/FasL mediated apoptosis in infected and chronically infected macrophages. Both mechanisms of apoptosis result in the death of the macrophage along with a portion of their intracellular Mtb. When adjacent, regulatory T cells can downregulate cytotoxic and IFN□ -secreting T cells, which leads to the loss of their cytokine-secreting and apoptosis-inducing abilities.

#### 2.2.4. Vascular blockage modeling

As cells in TB granulomas die, dead cellular debris accumulates into a necrotic core (caseum) that serves as a niche for bacterial survival but they cannot replicate; vascular sources are abnormal in caseum, leading to a hypoxic microenvironment^111^. We model vascular blockage depending on the amount of necrotic tissue (i.e., caseum) and cell density at the vascular source’s location. The blockage increases with higher necrotization and higher density. We model the level of necrotization as the ratio of the number of dead cells on that location to a maximum of dead cells allowed in each grid. Once the number of dead cells reaches the maximum value allowed, then we consider the grid as caseated, which means that the vascular source at that location is completely blocked, and cells can no longer move to that location.

#### 2.2.5. Mtb and bacterial proliferation

Mtb replicate based on their metabolic state, which differs based on their main niches: within-macrophages (intracellular Mtb), in the interstitial space between live cells (extracellular-replicating), and within caseum (extracellular-nonreplicating)^26^. Each Mtb agent in *GranSim* is assigned a metabolic mode based on their location: intracellular Mtb within macrophages, extracellular Mtb in the cellular region of granulomas and nonreplicating Mtb trapped in the caseum. Intra- and extracellular Mtb proliferate in *GranSim*, but immune cells and nonreplicating Mtb do not proliferate. The total rate of proliferation of intracellular Mtb depends on the adaptive immune response and decreases when adaptive immunity is effective. The rate of proliferation of extracellular Mtb depends on the caseation level and the number of Mtb within a grid compartment, with replication rate decreasing with increased caseation and Mtb burden.

#### 2.2.6. Chemokine-cytokine secretion

Similar to tumors, chemokines and cytokines are secreted within granulomas to activate, suppress, or chemoattract cells^26^. In *GranSim*, activated, infected and chronically infected macrophages secrete the chemokines CCL2, CCL5 and CXCL9. Unless they are downregulated by Tregs, Tcyts and Tgams are able to secrete cytokines—both Tcyts and Tgams secrete TNFα, while only Tgams cells secrete IFN□. Macrophages respond to the chemoattraction of CCL2 and CCL5, Tcyts to CCL5 and CXCL9, Tregs along CCL5, and Tgam along all CCL2, CCL5, CXCL9.

#### 2.2.7. Recruitment

Cytokines also inform the rate of recruitment of immune cells to the site of Mtb infection^110^. We recruit macrophages and T cells from vascular sources. The recruitment rates at each vascular source are dependent upon the concentrations of CCL2, CCL5, CXCL9, and TNF-α in the specified grid compartment similar to the description for *TumorSim*.

#### 2.2.8. Age-related cell death

As granulomas can persist for years^24^, we allow simulated cells to age and eventually die. During initialization and recruitment, the age of immune cells is randomly sampled between zero and their maximum age. Immune cells die when they reach their corresponding maximum ages.

### 2.3. Granuloma and tumor simulations

Though both *GranSim* and *TumorSim* share many immune components, the underlying biology suggests very different simulation endpoints. Unlike the immune-controlled growth of granulomas, aggressively-growing SCLC tumors continue to expand, often exceeding 3cm in diameter^112^. We expect that a virtual SCLC tumor without any immune resistance may exceed a 5mm x 5mm region within 15-30 days and overflow our simulated domain; for this reason, we simulate 2 weeks of tumor growth early after the tumor is established. By contrast, we simulate TB granulomas for 300 days to capture both early formation and late-stage maintenance.

To accurately understand how model outcomes depend on parameter values, we ensure that our parameter values give rise to each simulation well-represent the underlying biology. To ensure a thorough exploration of parameter space, we employ the Latin hypercube sampling (LHS) method^113^ that creates a stratified random sampling of a given parameter space^113,114^. For *TumorSim*, we generate 500 parameter sets using the parameter ranges described in Tables 2-8. For *GranSim*, we sample 500 parameter sets using parameter ranges calibrated in our previous work^66,67^. To address stochasticity inherent to the ABM (i.e., aleatory uncertainty) in both *TumorSim* and *GranSim*, we perform 3 replicate simulations for each parameter sample for a total of 1500 virtual SCLC tumors and 1500 virtual granulomas.

### 2.4. Classifying granuloma and tumor outcomes

Several common terms have been used to describe tumor status. Uncontrolled proliferation of cancer cells within tumors is frequently described as tumor progression^9,10,77^. Lack of progression is typically characterized by progression-free survival of the patient. Tumor regression is clinically defined as the disappearance of a tumor^115^. Tumors that neither progress nor regress are called dormant, whether that is due to a balance of pro- and anti-tumor effects, or a consequence of quiescent cells^116^. The vast majority of SCLC tumors progress, with few cases of regression ever reported^115,117^.

In *TumorSim*, we use simulated tumor volume to categorize simulated tumors into three analogous qualitative categories: progressing, dormant, or regressing. We compute tumor volume by counting the number of tumor-associated cells within the simulation. We then infer tumor radius by approximating the tumor area as a circle and estimating tumor volume by assuming a spherical tumor. We describe simulated tumors as progressing if volume increases near-exponentially, dormant if tumor volume plateaus, and regressing if the volume decreases or if all cancer cells die.

Similarly, common terms exist to describe TB granulomas. However, granuloma volume is driven by leukocyte count and the bulk of caseous necrosis, fibrosis, and calcification; macroscopic granulomas may be entirely sterile^118^. Rather, granuloma status is described based on their ability to control Mtb, including sterile, poorly-controlling Mtb, or well-controlling Mtb (although caseum is associated with poor control)^118^. Poorly-controlling granulomas are associated with the Mtb dissemination, causing disease severity and/or the foundation of new granulomas^24,26,118^.

In *GranSim*, we use total Mtb count to classify simulated granulomas into one of three states: sterilizing, controlling, and disseminating. Sterilizing granulomas are those that completely clear Mtb within 300 days. Controlling granulomas are those that maintain stable, low levels of Mtb after the adaptive immune system is primed. Disseminating granulomas are those that fail to control Mtb replication, characterized by a continued increase of Mtb throughout the simulation.

### 2.5. Uncertainty and sensitivity analyses

Complex models like hybrid ABMs are often nonlinear and have hundreds of inputs. Uncertainty and sensitivity analysis allow us to explore parameters that significantly impact outputs of interest. To uncover which dynamics drive heterogeneity in SCLC tumor and TB granuloma outcomes, we employ the partial rank correlation coefficient (PRCC) method, a robust sensitivity analysis method that quantifies nonlinear monotonic relationships^113^. Even small PRCC values may indicate significant correlations^113^, and so we compute *p-*values associated with each parameter adjusted for multiple comparisons via the Benjamini-Hochberg correction^119^. We use the PRCC method to analyze the 1500 virtual SCLC tumors and 1500 virtual granulomas generated previously.

## 3. Results

### 3.1. TumorSim and GranSim capture spatiotemporal features of SCLC tumors and TB granulomas

To validate *TumorSim*, we observe gross simulation output and ensure that the simulations reasonably reflect reality as compared to published data on lung lesions. In (Figure 3), we also present side-by-side comparisons between *TumorSim* and *GranSim* outputs to compare and contrast. In Figure 3A, we present a small lymphocyte-infiltrated SCLC tumor with an effective immune response as observed in human tumors^120^. Activated Tcyts are located near M1 macrophages and mostly at the tumor boundary as seen *in vivo*^*121*^. We see deactivated Tcyts near Tregs within the tumor. Most larger tumors had substantially fewer infiltrating leukocytes, growing essentially unchallenged by immune cells; such tumors are described elsewhere as “cold” and associated with poor outcomes^33,122^. While the majority of tumors continue to progress (Methods), a small number tend towards dormancy and regression (Figure 3C). While events of spontaneous SCLC regression are exceedingly rare clinically^115^, the small size of these simulated tumors (∼<200) are likely more comparable to the elimination stage of tumor development that cannot be observed *in vivo*^77,78^. As tumors grow, so do total numbers of overall tumor-associated T cells and macrophages (Figure 2C-G). By comparison, a majority of simulated TB granulomas are either controlling or sterilizing. Figure 3B shows a representative latent virtual granuloma, where the immune system controls but does not completely clear Mtb infection. Necrotic caseum accumulates in the center of the Mtb-infected lesion. Overall, we observe an early spike in Mtb count due to the delay in the adaptive immune response, whereafter granuloma volume, Mtb count, macrophage, and T cell count stabilizes for most granulomas (Figure 2H-L); this reflects the tendency for most granulomas to permit a persistent, latent Mtb infection state^24^. *GranSim* simulations also capture active TB disease dynamics, as over 40% of our simulated granulomas disseminate, with increasing Mtb counts throughout the simulation (Figure 2I). Supplemental Material S2 presents simulation outcomes over time for progressing tumors, dormant tumors, and regressing tumors; and non-controlling, controlling, and sterilizing granulomas; and Supplemental Videos S1-S3 show the development of the tumors in Figure 2A and supplemental Figure S2 (Supplement and videos located on our website: http://malthus.micro.med.umich.edu/TumorSim/).

**Figure 2:**
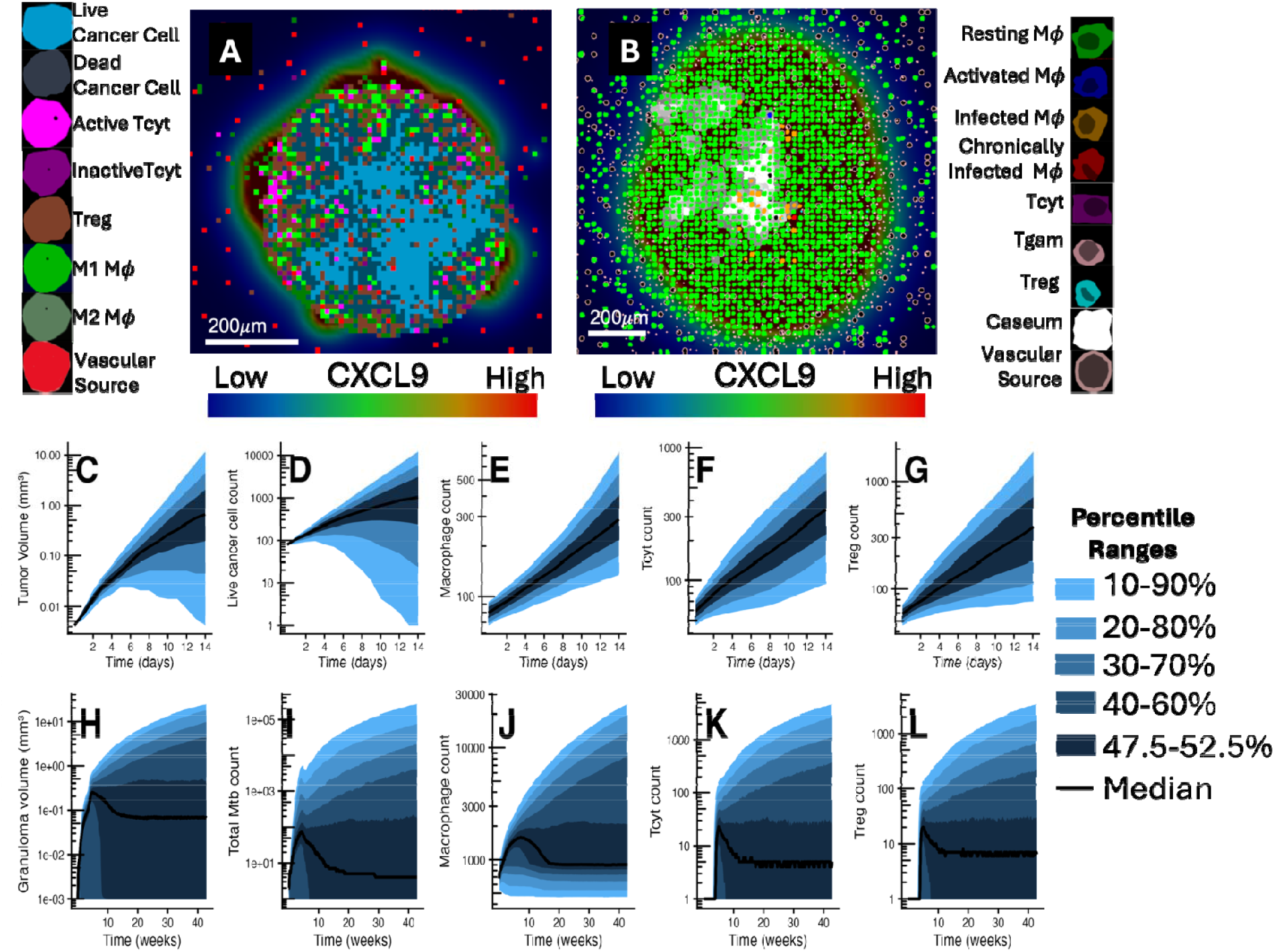
Spatial and Temporal *GranSim* and *TumorSim* model outputs for comparison and contrast. Both (A) *TumorSim* and (B) *GranSim* simulate spatial interactions of multiple cell types. Both scale ars are 200 microns. (A) A representative *TumorSim* SCLC tumor simulation of a lymphocyte-infiltrated tumor after 14 days of simulation. A mass of cancer cells (blue) is infiltrated by a collection of Tcyts and macrophages, while some Tcyts are being deactivated by Tregs We show relative levels of CXCL9 as a background heatmap. (B) A representative simulation of a non-sterile granuloma at day 300. Caseum has formed in the center of the infection site and is surrounded by macrophages as well as lymphocytes. We show relative levels of CXCL9 as a background heatmap. (A-B) We chose both simulations to demonstrate interaction of multiple cell types. (C-L) Each panel shows the median and percentile ranges for multiple temporal outcomes over time for both *TumorSim* (C-G) and *GranSim* (H-L). Outcomes include lesion volume, Mtb/Cancer cell counts, and leukocyte counts *GranSim* simulations are run for 300 days to capture both early- and late-state dynamics. *TumorSim* virtual tumors are simulated for two weeks, due to aggressive tumor growth.

**Figure 3:**
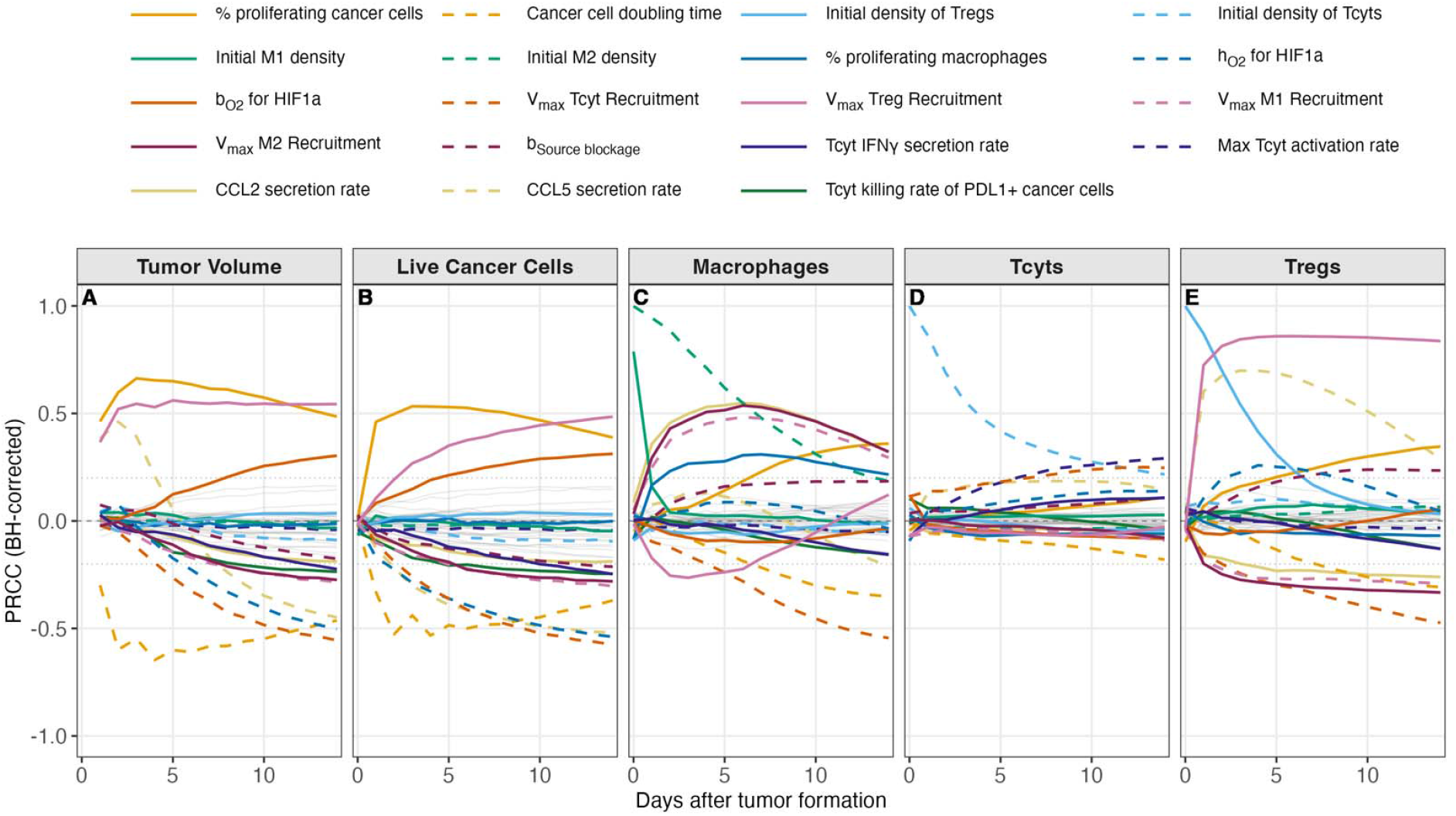
Parameters influencing *TumorSim* outcomes. To assess mechanisms that influence dynamics and outcomes of *TumorSim*, we present PRCC values over time for every varied *TumorSim* parameter (Methods). We have highlighted PRCC trajectories for those parameters that had significant correlation and |PRCC| > 0.2 for at least one outcome for one or more timepoints (p<0.05 after Benjamini-Hochberg multiple comparison correction^119^), while all other PRCC values are shown in gray. PRCC values presented capture the impact of parameters on the model outcomes: (A) tumor volume, (B) live cancer cell count, (C) number of macrophages, (D) Tcyt count, and (E) Treg count. Some C_50_ values were omitted from presentation for clarity when the associated V_max_/R_max_ parameters appeared as significant showing similar mechanistic impact.

### 3.2. TumorSim captures distinct roles of chemokines during tumor development

With the ability to mechanistically simulate 2 large sets of heterogeneous virtual granulomas and tumors, respectively, we analyze each model to determine which parameters (and thereby mechanisms) underpin the heterogeneity of outcomes observed during formation of the lesions. We measure partial rank correlation coefficients (PRCC; see methods) to capture how parameters that we vary in Tables 2-8 impacted five different tumor-scale outcomes (Figure 3). Throughout simulations, higher levels of Treg recruitment and faster cancer cell proliferation are associated with larger tumors. Beyond this, we notice a unique separation of early- and late-phase behavior of tumor growth that is driven by distinct mechanisms, occurring near the 1-week timepoint (Figure 3). Early during tumor growth, macrophage and Treg counts correlate with CCL2 and CCL5 secretion rates in addition to their initial counts. During this time, CCL5 positively correlates with tumor volume— likely due to the recruitment of Tregs. This correlation becomes negative later during tumor growth, around the time that Tcyt killing of PDL1+ becomes significantly negatively correlated with tumor volume. This later-phase shift also coincides with the time that Tcyt levels correlate with Tcyt activation probability - likely due to the importance of IFN□ production in T-cell recruitment via CXCL9 production by M1 Macrophages. While it is well-studied that a combination of CCL5 and CXCL9 perform a significant anti-tumor role^34,123^, our simulations suggest that CCL5 may serve a pro-tumor role early during tumor development by recruiting Tregs in the absence of CXCL9. It is during this later stage that PD1-Tcyt killing of PDL1+ cancer cells negatively correlates with tumor volume.

We also observe that macrophage populations negatively correlate with Treg recruitment, likely due to our assuming slower proliferation rates of M2 macrophages. We notice that as simulations progress, all cell populations correlate with faster cancer cell proliferation, likely due to higher net exposure of vascular sources to tumor-associated chemokines. Finally, we observe that as tumors develop, hypoxia-related parameters become more significant, with higher levels of HIF1α correlating with larger tumors, more cancer cells, larger Treg populations, and macrophage counts, likely due to increased Tcyt PD1 expression. Lastly, we note that no initial conditions of macrophages or T cells showed as significant to tumor outcomes, suggesting that long-term outcomes are dependent mostly on recruiting effects, which is consistent with development of adaptive immunity.

### 3.3. Granuloma outcomes are determined by a balance of pro- and anti-inflammatory regulators

From observing spatial and temporal dynamics it is clear there are similarities and differences between behaviors of SCLC and TB systems. To compare, we perform a parallel sensitivity analysis identifying what features control TB lesion outcomes in contrast to SCLC. As above, we apply the PRCC method to assess the impact of each mechanism on five *GranSim* outcomes analogous to *TumorSim* outcomes. We observe overlapping mechanisms driving bacterial burden and granuloma size, meaning granulomas with higher bacterial burden tend to have larger sizes. Concentration of Interleukin-10 (IL-10), a key anti-inflammatory cytokine, shows significant correlation with larger granulomas and higher levels of Mtb, as observed previously^124^. Similarly, lower rates of TNF-induced apoptosis, lower recruitment of Tgam, lower cognate Tgam ratio and higher recruitment of macrophages all lead to higher Mtb burden and larger granulomas during the course of infection, consistent with previous experimental and computational studies^29,125-128^. In addition, granuloma size is also sensitive to the thresholds of chemokines needed for macrophage recruitment in the beginning of the infection, such that higher thresholds lead to smaller granulomas due to decreased macrophage recruitment early on. Notably, by around 22 weeks, the drivers of these dynamics settle and we observe no change between drivers of granuloma volume between days 150 and 300, despite poorly-controlling granulomas continuing to increase in volume (Figure 2H).

## 4. Discussion

Understanding how cell-scale behavior influences macroscopic outcomes is key to understanding the pathology of SCLC and which potential treatments are most promising. Though many mechanisms about SCLC are known, effective treatments remain elusive. In this work we introduce a novel molecule-to-tissue scale model of a lung lesion, *TumorSim*, an agent-based model of SCLC tumors rooted in key cellular and cytokine players to probe which mechanisms principally drive variability in SCLC tumor outcomes.

We base *TumorSim* on the computational framework of *GranSim*, our hybrid ABM representing pulmonary TB granuloma formation^27-30^. Lung tumors and TB granulomas share multiple similarities, both in terms of clinical diagnosis^2,3^ and known drivers of the granuloma/tumor immune microenvironment^4^. Not surprising, data suggest that lung lesions are similar in many immune characteristics and even are mistaken for each other on lung examination^4^. Implementation of *TumorSim* utilizes similarities and differences describing immune-interaction mechanisms detailed in *GranSim* (Table 1), including our grid-based, lung-tissue simulation environment, molecular- and cell-scale programming of macrophages by T cells in various (anti-)inflammatory states, and an updated oxygen diffusion algorithm. *TumorSim*-specific components include PD1 and PDL1 interactions between T cells and cancer cells, descriptions of distinct macrophage states and hypoxia-driven vasculature blockage. We validated *TumorSim* by recapitulating multiple known drivers of tumor outcomes, including the relative interdependence of tumor outcomes on cytotoxic T-cell-to-macrophage interactions via CCL5 and CXCL9^34,123^, as well as the impact of cytotoxic T-cell activity on reduced tumor volume.

Examining *TumorSim* dynamics suggests that tumor dynamics early during tumor escape may be distinct from dynamics driving more mature tumors. Simulations predict that increased CCL5 serves a pro-tumor role during early tumor development by recruiting regulatory T cells before macrophages can mediate recruitment of cytotoxic T cells. During this early phase, Tregs protect the tumor before hypoxia-driven mechanisms become relevant. These could serve as targets for immune boosting treatments of tumors.

We observed several similarities and differences between SCLC tumors and TB granulomas. First, absolute numbers of all cell types—both pro-lesion and anti-lesion—appeared to scale with lesion size. Potent negative regulators (e.g., PD1/PDL1 in SCLC; IL-10 in TB; Tregs in both) had consistent impacts on lesion volume and cancer cell/Mtb count. By contrast, TB granulomas are much more effectively limited by the replicative niche of macrophages even when infection is not cleared; whereas non-dormant SCLC lack such constraints and can replicate indefinitely. This is reflected in SCLC proliferation rate correlating with tumor volume, while Mtb replication rate has little correlation with overall granuloma volume or Mtb count.

One goal of this study was to compare what we learn about both lesion types and their driving mechanisms for intervention purposes. Both *TumorSim* and *GranSim* show improved outcomes with increased cytotoxic activity: tumor volume negatively correlates with Tcyt and M1 macrophage recruitment (Figure 3), while granuloma Mtb count negatively correlate with Tgam, Tcyt, and antibiotic macrophage activity (Figure 4). Outcomes worsen with the action of regulatory T cells, as Tregs correlate with both increased tumor and granuloma volume.This suggests that targeted inhibition of Treg recruitment or activity could improve both SCLC and TB outcomes. We acknowledge that there is important nuance when considering immune and pharmacological interactions between SCLC and TB treatment. Previous (or present) Mtb infection substantially increases the chance of development of lung cancer^4^, yet immune checkpoint inhibitor treatment of lung cancer has been associated with reactivation of latent Mtb infection^129^. Still, given the prevalence of TB relapse post-antibiotic treatment^130^, it is plausible that many relapse-free TB patients could harbor granulomas after treatment. These granulomas may continue to reshape the lung immune environment via upregulation of local or systemically primed immunosuppressive lymphocytes, possibly resulting in increased Treg involvement during early tumor development.

**Figure 4:**
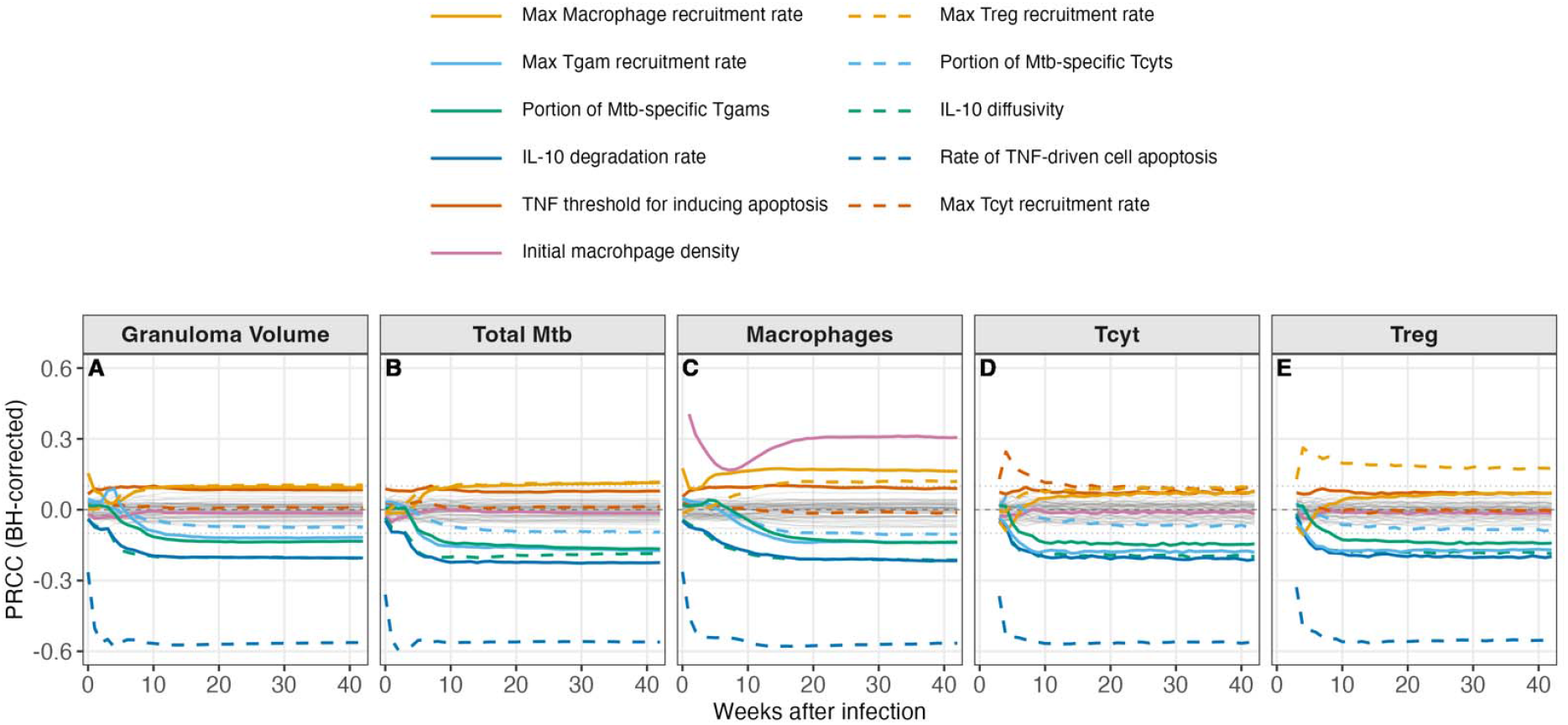
Parameters influencing *GranSim* outcomes. We present PRCC values over time for every varied *GranSim* parameter for several outcomes. We have highlighted PRCC trajectories for those parameters that had significant correlation for at least one outcome for at one or more timepoints (p<0.05 with Benjamini-Hochberg multiple comparison correction), while all other PRCC values are shown in gray. PRCC values presented capture the impact of parameters on the model outcomes: (A) granuloma volume, (B) total live Mtb count, (C) number of macrophages, (D) Tcyt count, and (E) Treg count.

Interestingly, we noted that in *TumorSim*, there was no significant correlation between initial levels of macrophages or T cells and tumor outcomes, suggesting that long-term outcomes are dependent mostly on recruiting effects, which is consistent with development of adaptive immunity. This may not be a true representation as 1) we are only simulating for 14 days and 2) tissue resident memory cells are known to be present in lungs and their role in the elimination phase of SCLC tumor development may contribute to variation in outcomes of people with SCLC or developing lung lesions^78^. Advancements in single-cell technology in exploring these early immune interactions may soon allow for more detailed simulations closer to the moment of malignant transformation^131^.

*TumorSim* is a first-generation step toward building a comprehensive model of lung cancer lesions. Though there is a rich and complex body of literature exploring many cell types and cytokines associated with SCLC, it is necessary to simplify the biology for computational tractability. Some biologically-distinct cell types have very similar cell-cell interactions at *TumorSim*’s level of biological resolution, e.g., myeloid-derived suppressor cells (MDSCs) and M2 macrophages both are recruited anti-inflammatory cells^9^. The action of neutrophils and natural killer cells have been included in the action of M1 macrophages and Tcyts, respectively. Some cell types are present only in small numbers (e.g., B cells)^12^. Further, some cells are more structural by nature (e.g., fibroblasts^12^), and are implicitly represented as part of the *TumorSim* grid as background lung tissue. Similarly, several continuous agents have been omitted for similar reasons, e.g., chemokines associated with omitted cell types. Finally, necrosis is a typical feature of SCLC, but is not always found in small samples comparable to those we simulate^132^. Future versions will explicitly represent these factors and others that are emerging as relevant in SCLC.

The modular structure of our hybrid ABM *TumorSim* will allow us to integrate emerging data for more detailed simulations. Further development will allow this platform to be the basis for hypothesis testing and exploring effects of treatment with chemotherapies and host-directed therapies. Detailed simulation of PD-1 blockade will allow us to investigate spatiotemporal effects and cell-cell interactions that underpin heterogeneous treatment responses that clinically define hot vs. cold tumors; we may also investigate how this correlates with immune-infiltration based notions of tumor ‘heat’^122^. *TumorSim* may provide valuable insights for treatment of SCLC or other immunologically-similar carcinomas^94^ which are the subject of our current work.

## Data availability

All data generated or analyzed during this study are included in this published article and are available upon request to the corresponding authors. Additional model details for generating simulations in this paper are available on our on-line supplements. For *TumorSim*, additional information is located at http://malthus.micro.med.umich.edu/TumorSim/, and for *GranSim*, additional details and parameter values are available at http://malthus.micro.med.umich.edu/GranSim/docs/gransimrules-v2.pdf.

## Acknowledgments

C.T.M. was supported by the Department of Microbiology and Immunology at the UM. This research was supported by the Advanced Cyberinfrastructure Coordination Ecosystem: Services & Support (ACCESS) program through allocations of hours of computer resources # MCB140228. We also thank the Center for Data-Driven Drug Development and Treatment Assessment (DATA; which supported M.B.). We thank Paul Wolberg for computational assistance and support.

## Notes

http://malthus.micro.med.umich.edu/TumorSim/

### Competing Interest Statement

The authors have declared no competing interest.

